# Lateralized Neural Entropy modulates with Grip Force during Precision Grasping

**DOI:** 10.1101/2023.05.07.539751

**Authors:** Nishant Rao, Andrew Paek, Jose L. Contreras-Vidal, Pranav J. Parikh

## Abstract

When holding a coffee mug filled to the brim, we strive to avoid spilling the coffee. This ability relies on the neural processes underlying the control of finger forces on a moment-to-moment basis. The brain activity lateralized to the contralateral hemisphere averaged over a trial and across the trials is known to be associated with the magnitude of grip force applied on an object. However, the mechanistic involvement of the variability in neural signals during grip force control remains unclear. In this study, we examined the dependence of neural variability over the frontal, central, and parietal regions assessed using noninvasive electroencephalography (EEG) on grip force magnitude during an isometric force control task. We hypothesized laterally specific modulation in EEG variability with higher magnitude of the grip force exerted during grip force control. We utilized an existing EEG dataset (64 channel) comprised of healthy young adults, who performed an isometric force control task while receiving visual feedback of the force applied. The force magnitude to be exerted on the instrumented object was cued to participants during the task, and varied pseudorandomly among 5, 10, and 15% of their maximum voluntary contraction (MVC) across the trials. We quantified neural variability via sample entropy (sequence-dependent measure) and standard deviation (sequence-independent measure) of the temporal EEG signal over the frontal, central, and parietal electrodes. The EEG sample entropy over the central electrodes showed lateralized, nonlinear, localized, modulation with force magnitude. Similar modulation was not observed over frontal or parietal EEG activity, nor for standard deviation in the EEG activity. Our findings highlight specificity in neural control of grip forces by demonstrating the modulation in sequence-dependent but not sequence-independent component of EEG variability. This modulation appeared to be lateralized, spatially constrained, and functionally dependent on the grip force magnitude. We discuss the relevance of these findings in scenarios where a finer precision is essential to enable grasp application, such as prosthesis and associated neural signal integration, and propose directions for future studies investigating the mechanistic role of neural entropy in grip force control.

## Introduction

Efficient grip force control is prominent in routine tasks such as holding a coffee mug that is filled to the brim. The seemingly simple task belies the underlying physiological complexity that is evident from the aberrant force control in pathological conditions e.g., stroke, Parkinson’s disease, cerebral palsy, and focal hand dystonia (Chu & Sanger, 2009; Fellows & Noth, 2003; Grafton, 2010; Lodha et al., 2013; Mishra et al., 2022; Olivier et al., 2007). Despite the ubiquitous nature of the task, neural mechanisms underlying grip force control remain elusive.

Activity within the frontal and parietal cortical regions is known to be associated with the magnitude of grip forces exerted on a hand-held object (Davare et al., 2011; Ehrsson, Fagergren, Jonsson, Westling, Johansson, et al., 2000; Ehrsson et al., 2003; Grafton, 2010; Kilner et al., 2003; Poon et al., 2013; Vaillancourt et al., 2003). During grasping, the fronto-parietal cortical activity is often lateralized to one hemisphere as a function of task phases and the processing requirements for grip force application (Davare et al., 2011; Ehrsson, Fagergren, Jonsson, Westling, Johansson, et al., 2000; Ehrsson et al., 2003; Grafton, 2010; Kilner et al., 2003; Poon et al., 2013; Vaillancourt et al., 2003). Notably, the role of cortical activity during force control has been typically assessed by the average neural signals (or changes in mean value). An alternative view suggests assessment of the task-dependent cortical dynamics by characterizing the fluctuations in neural activity, an aspect which is analytically undermined by averaging or pooling the activity either over the trial duration or across multiple trials (Aguirre et al., 1998; Birn, 2012; Grady & Garrett, 2018). Recent evidence highlights the critical role of neural variability from trial-to-trial in several behavioral paradigms such as perceptual matching, attentional cueing, face recognition, and delayed match-to-sample tasks (Garrett et al., 2010, 2011, 2014; Garrett, Kovacevic, et al., 2013; Garrett, Samanez-Larkin, et al., 2013). Importantly, the spatial pattern depicted by trial-to-trial neural variability during these tasks was more reliable and distinct from that revealed by averaging (or pooling) the cortical activity across trials (Garrett et al., 2010). These findings reinforced the notion that the brain signal variability is mechanistically relevant for behavior, beyond the role of average neural activity, and the former encompasses spatially distinct and functionally orthogonal neural mechanisms from that identified by the latter (Garrett et al., 2011, 2014; Garrett, Kovacevic, et al., 2013; McIntosh et al., 2010). Concomitant with this framework, we previously showed a systematic modulation in trial-to-trial variability in corticospinal activity which occurred above and beyond the changes explained by the averaged activity across the trials, prior to grasping an object (Rao & Parikh, 2019a). This work supported the notion that the neural variability might inform not only cognitive functions involving face matching or attentional cueing, but also the finer sensorimotor behavior involving grip force control during grasping.

The capability to record neural signals noninvasively at an enhanced temporal resolution using electroencephalography (EEG) has also enabled the investigation of neural variations within a trial. For instance, variability in the cortical activity quantified using sample entropy of the EEG activity showed systematic increase in the signal complexity with developmental maturation (Mcintosh et al., 2008). Importantly, sample entropy provides a direct assessment of the sequential structure within the neural signal thereby, complementing the frequency-based spectral assessment which is sequence-independent. While our previous study showed spectral changes in EEG activity during a grasping task, much less is known about the sequence-dependent changes in neural variability within a trial during grip force control (Paek et al., 2019).

Based on the spatially lateralized fronto-parietal cortical processing for grip force control (Paek et al., 2019; Perez & Cohen, 2009; Rao & Parikh, 2017, 2019a), we hypothesized that the variability in EEG activity recorded over the fronto-parietal region will be lateralized and modulated with the increase in grip force magnitude. We characterized EEG variability using standard deviation (SD) and sample entropy (sE). While SD quantifies deviation in cortical activity around its mean, sE is known to quantify the degree of regularity in a signal such that higher entropy indicates lesser regularity (and increased complexity) whereas lower entropy indicates higher regularity (and lesser complexity) (Lodha & Christou, 2017; Mcintosh et al., 2008; Rao, Skinner, et al., 2019; Shah-Basak et al., 2020). Notably, SD is sensitive to the scaling of signal amplitude but invariant to its underlying sequence, whereas sE is sensitive to the sequential structure of the signal but invariant to the constant scaling of the signal amplitude – making these measures complementary for quantifying neural variability (Garrett, Samanez-Larkin, et al., 2013; Grady & Garrett, 2014; Shah-Basak et al., 2020).

## Methods

### Participants

Eleven healthy young naïve, right-handed adults (three females, eight males; age range: 18-35 years) participated in the study (Oldfield 1971). The current study has been performed based on data collected at Arizona State University, and reported in our previous article (Paek et al., 2019). Subjects provided written informed consent in accordance with the protocol approved by the Institutional Review Board at the Arizona State University. We used data from 8 participants who performed the task using the same experimental protocol.

### Grip device

A customized instrumented grip device was used for this study. Two force sensors (six-dimensional force/torque transducers; Nano-25, ATI Industrial Automation, Garner, NC) were instrumented on each graspable side of the grip device to independently record force exerted on each graspable surface (**Figure 1**). A 12-bit analog to digital converter board (sampling frequency 1 kHz, PCI-6225, National Instruments, Austin, TX, United States) was used to acquire the force data.

**Figure 1:**
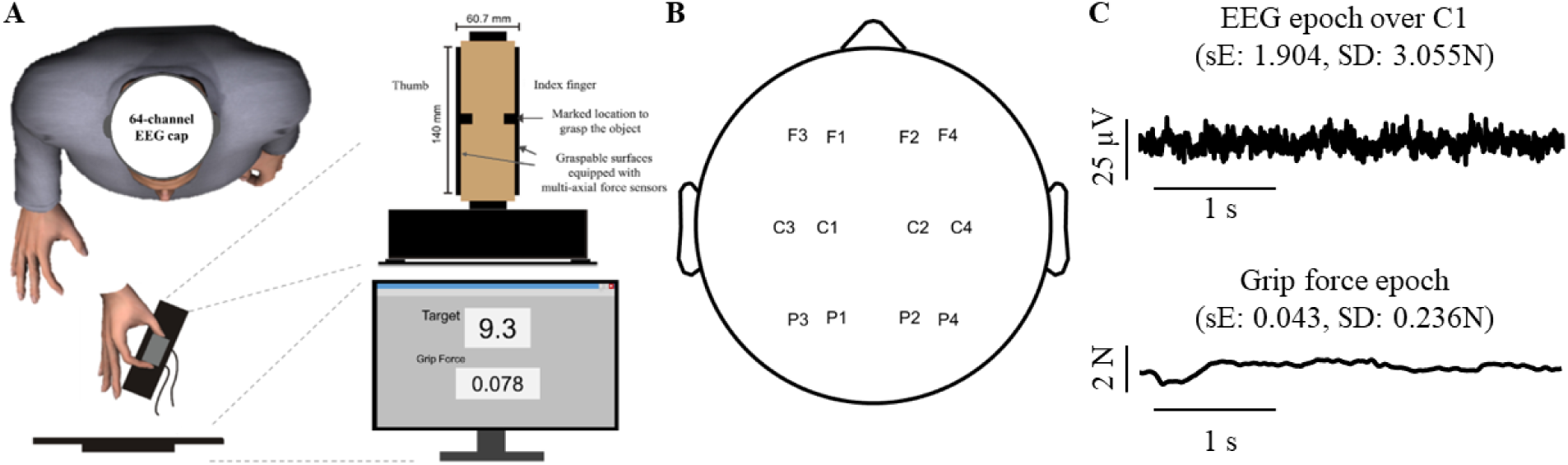
(A) Experimental setup showing a subject with EEG cap grasping the grip device while the applied grip and target forces are displayed on a screen; (B) Schematic of the ROI among the 64-channel EEG locations; (C) A single trial epoch from EEG over C1 (*top*) and grip force (*bottom*) data at 15% maximum voluntary force of a representative subject. sE and SD denote sample entropy and standard deviation, respectively.

### Electroencephalography

We used 64 channel standardized 10-20 EEG system (Goel et al., 2019; Luu et al., 2016) to assess the cortical activity of the participants as they performed the grip force task. The EEG system (BrainAmps DC amplifiers, Brain Products GLMB; sampling frequency 1 kHz for each electrode) consisted of one ground and one reference electrodes (placed on the subject’s earlobes) and four of the 64 channels dedicated to the measurement of electrooculography (placed near eyes and the temple) to get an improved estimate of eye movements for subsequent artifact removal (Paek et al., 2019). To ensure proper contact of the electrode with the scalp skin, the electrodes were supplied with saline gel.

### Experimental task

During the experimental sessions, participants were seated comfortably and instructed to perform an isometric force production task using their right hand. The customized object to be grasped was placed on a table in front of the chair (∼30cm away from the subject). Maximum voluntary contraction (MVC) for each participant was estimated as the maximal grip force exerted by the participant on the grip device when grasping only using their index finger and thumb (Lukos et al., 2013; Paek et al., 2019; Rao, Chen, et al., 2019; Rao & Parikh, 2017). For every participant, the experimenters performed three such repetitions to determine a consistent MVC (Paek et al., 2019; Rao & Parikh, 2019a). Before each trial, the subject was asked to bring their hand near the graspable surfaces of the object and cued to exert 5%, 10%, or 15% of their maximum voluntary contraction (MVC). The participants were also instructed to maintain this force level for 8s following which they were cued to stop exerting the force. The target force to be exerted as well as the actual grip force exerted by the subjects was displayed for the entire duration of the trial (**Figure 1**). At least hundred trials were recorded for each subject during the session resulting into ∼33 trials per force level. We recorded EEG activity (**Figure 1**) while subjects performed the trials.

### EEG preprocessing

We preprocessed the EEG data using steps formulated in (Delorme & Makeig, 2004; Paek et al., 2019). First, we re-referenced the sensor activity to that recorded from the mastoids (TP7, TP8) followed by removal of ocular artifacts using the H-infinity adaptive filter. The mastoid-based re-referencing primarily takes into the account any changes in sensors’ activity due to temporary shifts in ground and/or reference electrodes. Undertaking this step before the removal of ocular artifacts is essential to prevent loss of relevant information as H-infinity is sensitive to sudden changes in signal amplitude. Next, the possibility of a scalar drift (usually ∼0 Hz activity) in the signals was addressed by removal of any residual offset. This step was conducted by application of a high-pass filter at 0.1 Hz (zero-phase, 4_th_ order Butterworth filter). Notch filters were subsequently used to remove the power line noise and associated harmonics at 60, 120, and 180 Hz. Post line noise removal, it was important to detect changes in head movements and remove its influence from the neural signal using the artifact subspace reconstruction (ASR). Next, we applied independent component analysis (ICA) to remove the artifacts associated with muscle activity. Detailed description about these procedural steps is provided below.

After re-referencing the sensor activity with that in mastoids, the H-infinity adaptive filter was applied to remove the ocular artifacts (Kilicarslan et al., 2016). The electrooculography (EOG activity) was recorded by placing two sensors below the subjects’ right and the left eyes (IO1 and IO2 sensor) as well as near their temples (LO1 and LO2). Vertical EOG reference was computed by averaging the difference between EOG over IO1 and EEG over Fp1 (corresponding frontal activity), and that between EOG from IO2 and EEG over Fp2, as described below:

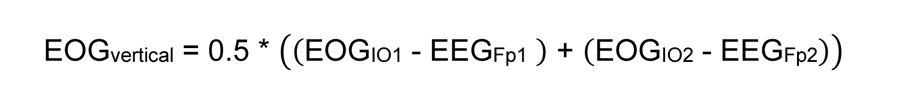

where EOG_vertical_ is the vertical reference, EOG_IO1_ and EOG_IO2_ represent EOG below the left and right eyes respectively, whereas EEG_Fp1_ and EEG_Fp2_ represent EEG corresponding to frontal activity from the left and the right side, respectively.

The horizontal EOG reference was computed by taking a difference between the EOG near the temples (LO1 and LO2) as described below:

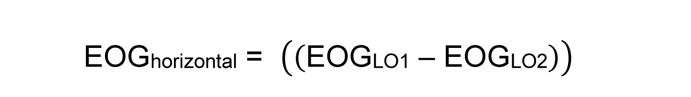

Where EOG_horizontal_ represents the horizontal reference for EOG, EOG_LO1_ and EOG_LO2_ represent EOG over the left and the right temples, respectively. After this computation, the H-infinity filter estimates the ocular artifacts using vertical and horizontal EOG references. As designed by Kilicarslan and colleagues and previously reported (Paek et al., 2019), the H-infinity filter is sensitive to the parameters *γ* and *q.* Based on these parameters, the filter adaptively estimates the output measures and isolates the ocular artifacts from the rest of the EEG activity. The parameters *γ* and *q* were optimized to 1.5 and 10^-20^ (determined using trial-and-error (Paek et al., 2019) to balance removal of the ocular artifact while preserving the integrity of EEG activity.

### Artifact Subspace Reconstruction

The H-infinity filtering was followed by removal of low-frequency offset (high-pass filter at 0.1 Hz) and power-line noise for 60, 120, and 180 Hz. The next step involved removal of brief head movements leading to aberrant EEG activity. This could be accomplished by the artifact subspace reconstruction (ASR) using *clean_asr* function in the MATLAB-based *EEGlab* toolbox. The ASR first identifies instances with unusual power spectrum (>10 standard deviations) in windows of 500 samples and step size of 250. This data is set to be reconstructed. Subsequently, the function generates epochs of 1-second-long data where no more than 3 EEG channels (and 2 for the pilot data) exceed power spectrum beyond -3.5 and 5.5 standard deviation compared to a robust EEG distribution. The ASR procedure estimates the robust distribution from each channel where the clean data is to be reconstructed based on truncated Gaussian distribution. Finally, the function reconstructs cleaner epochs for the instances identified as artifacts associated with sudden head movements, to be followed by removal of EMG artifacts using the ICA.

### Independent component analysis (ICA)

ICA was mainly applied to eliminate artifacts associated with muscle and forehead movements. First, a surrogate data stream was high pass filtered at 2 Hz to obtain a clearer estimate of scalp projections. This stream of data was run through the ICA cleaning procedure which identified components contributing to observed variance. These components were individually identified and removed if they (a) contained spectral power concentrated in the low frequency range within the frontal regions (characteristics of forehead movement) or (b) 50% power within the frequency range of 30 to 200 Hz with components localized to the peripheral scalp regions (typical attributes of muscle artifacts). An average of ∼35 components were identified and removed from the isometric force production task across subjects, and the resulting components representing cleaner EEG activity were utilized for subsequent analysis. The ICA-based approach has been consistent with that used in our previous studies (Kilicarslan et al. 2016; Goel et al. 2019).

### Data analysis

Based on our previous findings that highlighted the role of frontal, central, and parietal cortical activity during grip force control, we defined our region of interest (ROI) within the contralateral (left) frontal and parietal electrodes (i.e., F1, F3, C1, C3, P1 and P3) and their ipsilateral counterparts (i.e., F2, F4, C2, C4, P2 and P4). To test our hypothesis whether neural variability within the frontal and parietal regions was lateralized and systematically modulated with grip force magnitude, we computed the standard deviation and sample entropy (SD and sE respectively) in neural activity over the fronto-parietal brain regions (i.e., ROI defined above) using sensor-based activation. We focused our analysis on the variability (SD and sE) in time-domain EEG activity using a 1.6s long hold phase epoch. A computerized script as well as visual inspection were used to remove the initial ramp phase (de Freitas & Lima, 2013; Feeney et al., 2018). Parameters for sE (m=2, r=0.2) were consistent with previous reports (Svendsen & Madeleine, 2010; Vieluf et al., 2015).

Grip force data was assessed by first applying the zero-phase lag, fourth order, low-pass Butterworth filter with cutoff frequency of 14Hz (Flanagan & Beltzner, 2000; Rao & Parikh, 2019a) followed by computing mean, SD, sE, and coefficient of variation (CV = SD/mean) of grip force in the epoch defined earlier. Notably, SD and CV indicated amplitude-dependent and amplitude-normalized components of grip force variability. As SD and CV quantify sequence-independent variations, we also included sE to quantify the sequence-dependent component of grip force variability. We assessed changes in these variables with force magnitude using repeated measures analysis of variance (rmAnova, α=0.05) with *Force* (5, 10, and 15% MVC) as within-subjects factor. To assess the modulation in EEG variability across both hemispheres and with different force levels, we performed rmANOVA (α=0.05) on sE as well as SD of EEG activity recorded over the ROI including within-subjects factors such as *Laterality* (Left hemisphere, Right hemisphere), *Channels* (4 electrodes for each of the central (C1, C2, C3, C4), frontal (F1, F2, F3, F4), and parietal (P1, P2, P3, P4) topography in ROI), and *Force* (5, 10, 15% MVC). Notably, comparison between channels closer to the midline (e.g., C1 from left and C2 from right hemisphere) with those more lateral from the midline (e.g., C3 from left and C4 from right hemisphere) allowed us to assess the differences in EEG activity based on laterality and topographical proximity to the midline. We applied Huynh-Feldt corrections when the sphericity assumption was violated. Post hoc comparisons were performed using Tukey’s method. Statistical analyses were performed using SPSS software version 25.0 (IBM, USA).

## Results

We first provide findings related to the grip force execution during the epoch of interest, followed by changes in fronto-parietal EEG activity. The primary motive for this study was to characterize the modulation in neural variability with lateralized activation, channel topography, and the magnitude of grip force exerted on the object. We quantified the temporal structure of EEG variability (sequence-dependent) by computing the sample entropy of the EEG signal and assessed the sequence-independent changes by SD in the signal over the ROI.

### Modulation of grip force behavior with force magnitude

All subjects modulated their grip force as per the instructions and feedback provided (main effect of Force: F_(1.003,7.024)_=145.955, p<0.001, η_p2_=0.954; post hoc Tukey’s test for 5 vs. 10, 10 vs 15, and 5 vs. 15% MVC: all adjusted p<0.0001; **Figure 2**). The exertion of higher grip force magnitude also accompanied an increase in standard deviation in the grip force (main effect of Force: F_(2,14)_=23.305, p<0.001, η_p2_=0.769; post hoc Tukey’s test for 5 vs. 10% MVC: adjusted p=0.014, 10 vs. 15% MVC: adjusted p=0.025, 5 vs. 15% MVC: adjusted p=0.001). CV of grip force reduced with increase in magnitude of grip force (main effect of Force: F_(2,14)_=10.434, p=0.002, η_p2_=0.598) from 5 to 10% MVC (post hoc Tukey’s test: adjusted p=0.012) and from 5 to 15% MVC (post hoc Tukey’s test: adjusted p=0.009) but not from 10 to 15% MVC (post hoc Tukey’s test: adjusted p=0.640). The sample entropy in grip force reduced with increasing magnitude of grip force (main effect of Force: F_(2,14)_=9.923, p=0.002, η_p2_=0.586) especially from 10 to 15% MVC (post hoc Tukey’s test: adjusted p=0.018) and from 5 to 15% MVC (post hoc Tukey’s test: adjusted p=0.019), but not from 5 to 10% MVC (post hoc Tukey’s test: adjusted p=0.143).

**Figure 2:**
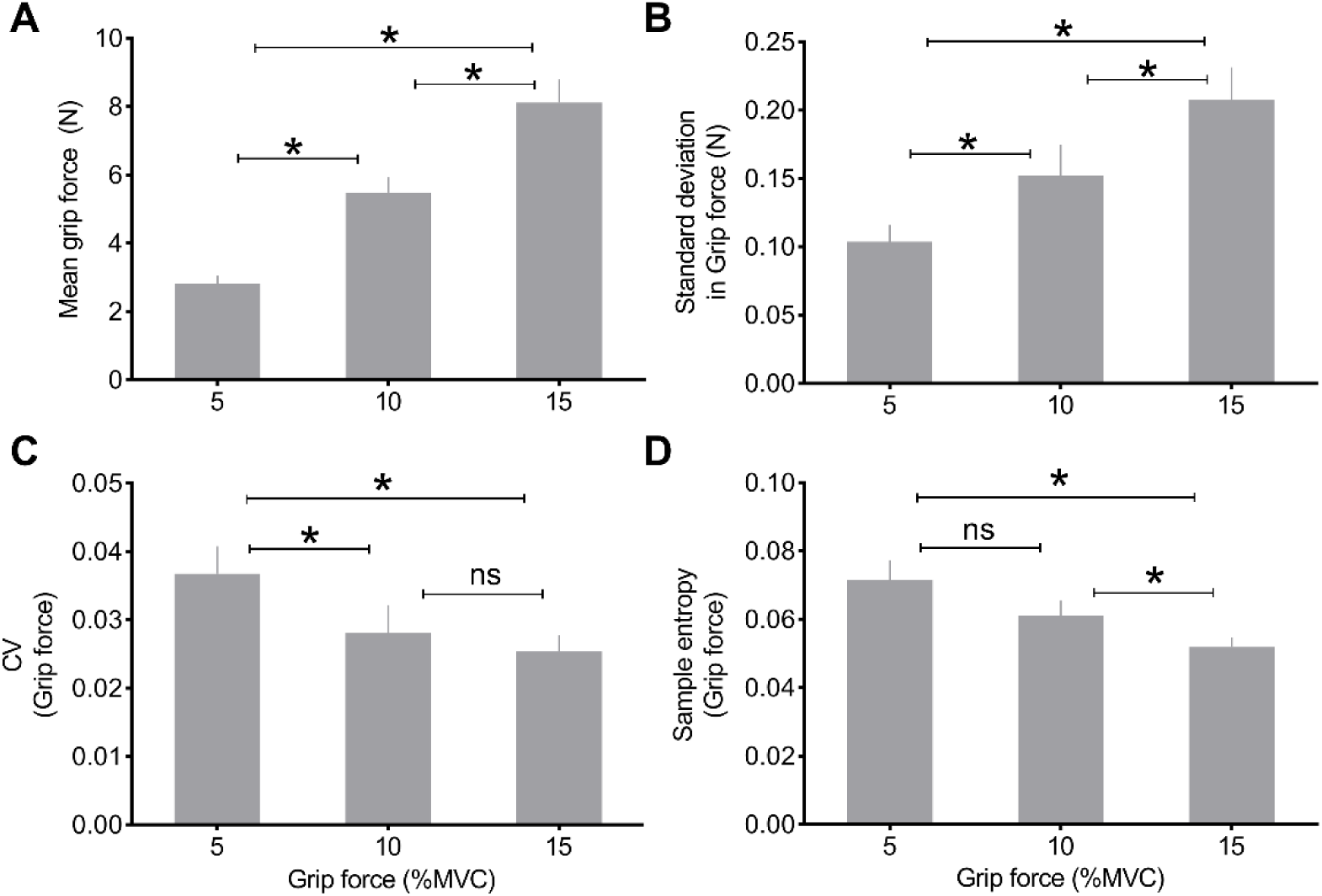
Changes in (A) mean, (B) SD, (C) CV, and (D) sE in grip force with force magnitude. Bars and error bars indicate mean and standard error respectively, asterisk indicates adjusted p<0.05 and ns indicates no statistical significance

### Sample entropy in EEG signals: Central electrodes

Sample entropy (sE) in EEG signals recorded over the central electrodes modulated based on the laterality and force magnitude (Laterality × Force interaction: F_(2,14)_=4.197, p=0.037, η_p2_=0.375; **Figure 3**). Post hoc comparisons showed statistically significant lateralized modulation when comparing C1 (left) vs. C2 (right) electrode activity from 10% to 15% MVC (post hoc Tukey’s test: adjusted p=0.048), but not from 5 to 10% MVC (post hoc Tukey’s test: adjusted p=0.687) and from 5 to 15% MVC (post hoc Tukey’s test: adjusted p=0.687). C3 versus C4 electrode activity showed no modulation in EEG activity with force magnitude (post hoc Tukey’s test: all adjusted p>0.246). Entropy did not alter based on the topographical proximity of the central electrodes from the midline (no Laterality × Channel interaction: F_(1,7)_=0.969, p=0.358; no Channel × Force interaction: F_(2,14)_=0.141, p=0.870; no main effect of channel: F_(1,7)_=4.667, p=0.068).

**Figure 3:**
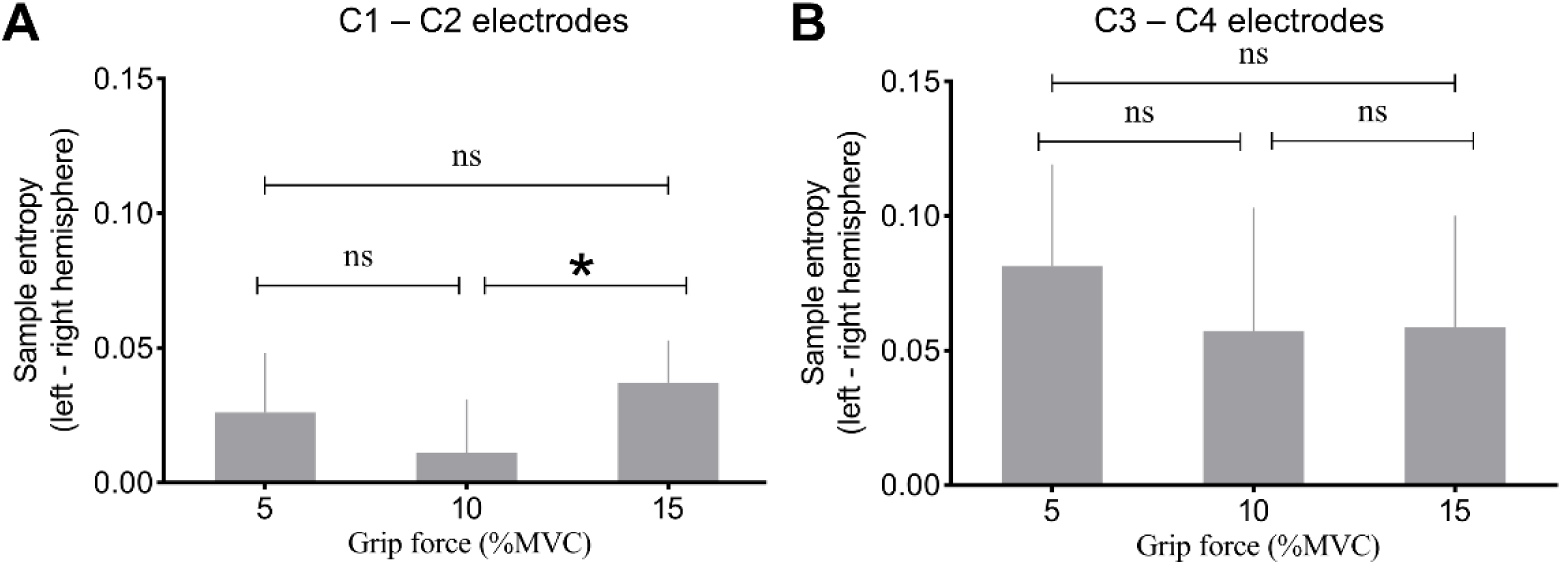
Lateralized entropy for three grip force magnitudes in EEG activity over (A) C1 minus C2 electrodes, and (B) C3 minus C4 electrodes; bars and error bars indicate mean and standard error respectively; asterisk indicates adjusted p<0.05 and *ns* indicates no statistical significance

### Sample entropy in EEG signals: Frontal and Parietal electrodes

sE in EEG activity within the frontal electrodes showed no modulation with laterality or force magnitude (no Laterality × Force interaction: F_(2,14)_=1.361, p=0.288; no main effect of Laterality: F_(1,7)_=0.014, p=0.910; no main effect of Force: F_(2,14)_=0.578, p=0.574). The frontal EEG activity showed no modulation with channel and force magnitude (Channel × Force interaction: F_(2,14)_=4.546, p=0.030, η_p2_=0.394; post hoc Tukey’s test: all adjusted p>0.400; no Laterality × Channel interaction: F_(1,7)_=0.228, p=0.648; no main effect of channel: F_(1,7)_=0.185, p=0.680; **Figure 4**).

**Figure 4:**
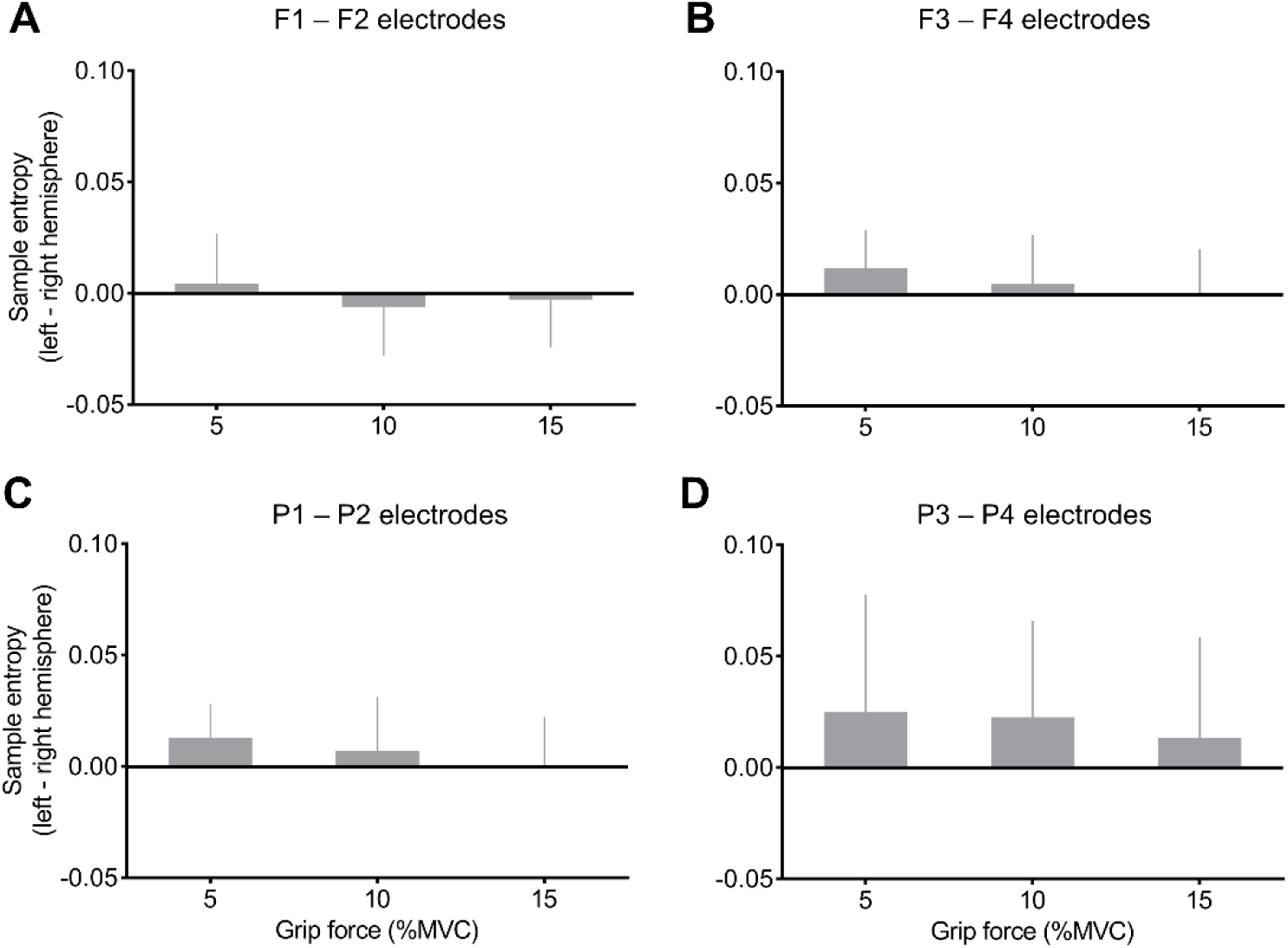
No modulation in sample entropy in EEG activity over the frontal and posterior parietal regions with brain laterality, nor with force magnitudes for (A) F1 minus F2, (B) F3 minus F4, (C) P1 minus P2, and (D) P3 minus P4 electrodes; bars and error bars indicate mean and standard error respectively.

Similarly, EEG activity in parietal electrodes showed no modulation with laterality, force, or channel (no Laterality × Force interaction: F_(2,14)_=0.436, p=0.655; no main effect of Laterality: F_(1,7)_=0.260, p=0.626; no main effect of Force: F_(2,14)_=0.701, p=0.513; main effect of channel: F_(1,7)_=10.318, p=0.015, η_p2_=0.596; post hoc Tukey’s test: all adjusted p>0.740; no Channel × Force interaction: F_(2,14)_=0.007, p=0.993; no Laterality × Channel interaction: F_(1,7)_=0.082, p=0.783; **Figure 4**).

### No modulation in SD in EEG activity with laterality, channel topography, and force magnitude

The standard deviation (SD) in EEG activity over central electrodes altered with laterality and force magnitude (Laterality × Force interaction: F_(2,14)_=4.358, p=0.034, η_p2_=0.384; no main effect of Laterality: F_(1,7)_=2.680, p=0.146; no main effect of Force: F_(2,14)_=1.037, p=0.380). However, none of the posthoc comparisons were significant (Tukey’s test for C1 vs. C2 electrodes activity: all adjusted p>0.071; for C3 vs. C4 electrodes: all adjusted p>0.410). No other contrast detected changes in EEG SD over central electrodes (no main effect of channels, nor associated interaction effects: all F<1.568, all p>0.250).

SD in EEG activity over frontal electrodes did not alter with channel topography (main effect of Channels: F_(1,7)_=19.701, p=0.003, η_p2_=0.738; post hoc Tukey’s test for F1 vs. F3, and F2 vs. F4 electrode activity: all adjusted p>0.990). No other contrast showed modulation in EEG SD from the frontal electrodes (no main effect, nor interaction effect: all F<2.440, all p>0.130). Similarly, EEG activity recorded over the parietal electrodes showed no change with channel topography (main effect of Channels: F_(1,7)_=12.891, p=0.009, η_p2_=0.648; post hoc Tukey’s test for P1 vs. P3, and P2 vs. P4 electrode activity: all adjusted p>0.850). Other contrasts failed to show modulation with laterality, force, or channel topography (no other main effects, nor interaction: all F<1.700, all p>0.234).

## Discussion

We investigated the modulation of cortical variability measured using EEG during an isometric grip force production task. We found lateralized, force magnitude-dependent modulation of sample entropy in electrical activity recorded over the central electrodes. Interestingly, this modulation appeared non-linear in nature. In contrast, similar modulation was not observed in the EEG activity recorded over the frontal and parietal regions. These findings indicate spatially constrained changes in the regularity of EEG signals during the force production task at the sensor-space resolution. On the other hand, assessment of sequence-independent component of neural variability (via SD) showed no modulation with laterality, channel topography, or force magnitude. We discuss these findings in the context of fronto-parietal cortical activity underlying grip force control and their relevance in the development of noninvasive brain-machine interfaces.

### Lateralized modulation in EEG variability during grip force control

Neural variability has been previously studied in the context of cognitive tasks involving decision making, face-matching, or attentional cueing paradigms (Mcintosh et al. 2008; Garrett et al. 2011b, a, 2014). In the current study, we highlight lateralized and force-magnitude dependent modulation in EEG variability over the central electrodes during a sensorimotor force production task in healthy young individuals. Prior neuroimaging work focusing on averaged neural activity has shown hemisphere-specific activation in sensorimotor and premotor cortical activity during force control tasks (Ehrsson et al. 2001, 2003; Poon et al. 2012, 2013). For instance, activation in contralateral primary somatosensory, motor, dorsal and ventral premotor cortices (S1, M1, PMd, and PMv respectively) has been shown to encode kinetic and kinematic representation in nonhuman primates (An et al., 2018; Atique & Francis, 2021). Other studies have observed similar spatial activation during the exertion of grip force in humans using fMRI as well as EEG (Ehrsson et al. 2003; Poon et al. 2012, 2013). Consequently, involvement of contralateral S1, M1, PMd, and PMv during the force control task could underlie the increase in EEG variability over central contralateral versus ipsilateral hemisphere. Studies probing mechanistic contribution of these regions highlight the processing of visual information specific to grip force scaling within contralateral M1 and PMd cortices (Davare et al. 2006, 2011; Olivier et al. 2007; van Polanen and Davare 2015). The integration of visuomotor information across M1 and PMd is known to be facilitated via the reciprocal tracts and phase-locked oscillations during the motor tasks (Atique & Francis, 2021; Churchland et al., 2006; Hendrix et al., 2009). Given the sensitivity of entropy to sequential interactions within the EEG signal, it is likely that the observed lateralized changes in entropy in electrical activity over central channels could represent the phase-dependent information processing within and between left M1 and PMd.

### Influence of cognitive and sensorimotor processes on neural variability

The lateralized changes in EEG variability over central electrodes in sE with force magnitude were non-linear in nature. An earlier work has reported a non-linear association between blood oxygen-level dependent (BOLD) signal over motor and non-motor areas and grip force magnitude (Alahmadi et al., 2016). A non-linear modulation in neural activation during task demands could be arguably traced to neurophysiological and cognitive sources (Alahmadi et al., 2016). The changes in attentional levels during production of grip forces at different levels under visual feedback might engage motor cortical regions that respond non-linearly as well as regions that respond linearly with varying grip force (Binkofski et al. 2002; Alahmadi et al. 2016. It is possible that the non-linear nature of modulation in EEG variability with force levels observed in our study might represent activity of motor cortical regions that are sensitive to attention levels and respond non-linearly to increasing grip force magnitude. Consistent with this notion, a recent study assessing intracortical activity recorded over M1 via implanted multiunit electrodes in patients with tetraplegia showed non-linear influence of volitional state (imagined, observed, or attempted movement) on grip force representation in M1 neural activity (Rastogi et al., 2020). Moreover, the authors found that neural activity encompassed by the implanted electrodes was particularly sensitive to the volitional states, while showing a subset of this activity also sensitive to the non-linear interaction of volitional state and force levels (Rastogi et al., 2020). As we observed reduction in lateralized neural variability over central electrodes at 10% MVC, it is possible that a combination of the cognitive and force level representation in the neural variability could be influenced by ’slacking’ (i.e., repeating drifts) in grip force, a phenomenon reported to exhibit a relationship with force magnitude that is linear for forces below 10% MVC, and non-linear for forces greater than 10% MVC (Smith et al., 2018).

### Applicability of neural variability to brain-computer interfaces (BCI)

Our study provides experimental evidence to the theoretical framework supporting mechanistic role of neural variability to grip force control. When considered alongside previous studies probing neural variability to understand cognition, we highlight neural variability as a potential feature to aid in the development of (non)invasive BCI applications for clinical rehabilitation. An important problem in the BCI literature has been to decode the motor intent to facilitate successful execution of action for patients with sensorimotor deficits e.g., stroke, Parkinson’s disease, tetraplegia, or spinal cord injury to name a few (An et al., 2018; Atique & Francis, 2021; Bhagat et al., 2016; Bradberry et al., 2009; Kumar, Das, et al., 2016; Kumar, Verma, et al., 2016; Luu et al., 2017; Moore et al., 2020; Paek et al., 2019; Park et al., 2021). Studies focusing on this problem have shown feasibility of decoding the motor intent signals from cortical recordings obtained invasively from nonhuman primates (An et al., 2018; Atique & Francis, 2021) as well as from semi-invasive electrocorticography in humans (Sanchez et al., 2008; Schalk et al., 2007). Importantly, our previous work showed success in decoding the motor intent from noninvasive EEG signals recorded from healthy young individuals, comparable to the extent of decoding the motor intent from invasive approaches (Bradberry et al. 2009, 2010). Thus, the potential of incorporating noninvasive signals in BCI applications for characterizing sensorimotor intent has been established and of considerable interest (Bradberry et al., 2010; Paek et al., 2019; Park et al., 2021; Yadav et al., 2019). Our findings pertaining to the neural variability within a trial (current study) as well as across the trials (Rao & Parikh, 2019a) support the notion that neural variability could provide additional insights into neural computations prior to, and during movement execution, and may enhance the decoding of motor intent for BCI applications.

With BCI systems equipped to execute the reaching movement using the neural signals, an important challenge faced by the current systems has been to incorporate specialized mechanisms enabling dexterous manipulation of the objects (An et al., 2018; Atique & Francis, 2021; Bhagat et al., 2016; Luu et al., 2017; Moore et al., 2020; Paek et al., 2019). Our recent study investigating reconstruction of grip force trajectories across range of force magnitudes using noninvasive EEG signals showed that the spectral power in theta (4-8 Hz), alpha (8-12 Hz) and beta bands (14-30 Hz) over the parietal areas was strongly correlated with grip force trajectories (Paek et al., 2019). On the contrary, spectral power in gamma bands (>30 Hz) over the frontal areas showed stronger correlations with the force trajectories (Paek et al., 2019). As our current findings depict non-linear modulation in EEG variability with grip force magnitude that was spatially specific to the central electrode locations but not the frontal and parietal electrode sites, we consider neural variability as a complementary measure to characterize grip force trajectories. Consequently, future studies could consider incorporating EEG signal entropy feature in addition to the currently known features (e.g., spectral power and voltage potentials) for reconstructing the grip force trajectories. While it is one of the strengths of the current study to highlight higher-order, neural entropy-driven dynamics during grip force control via sensor-space data for noninvasive BCI applications, it should be noted that the sensor-space resolution could be a limiting factor in determining the spatially specific activity within the selected frontal, central, and parietal electrodes. We acknowledge that increasing the spatial resolution could further advance our understanding of the finer interactions among neural ensembles. Collectively, our findings highlight the complementary nature of neural variability that is sequence-dependent, spatially specific, and functionally sensitive to the grip force task demands.

Findings from the current study also prompt further investigation into the causal role of the frontal and parietal cortical circuits in force dynamics (Olivier et al., 2007; Parikh & Cole, 2015; Rao, Mehta, et al., 2020; Rao & Parikh, 2019b; Young et al., 2020). Within the frontal lobe, dorsolateral prefrontal cortex, supplementary motor area, dorsal and ventral premotor, and the primary motor cortices are known to be involved in processing behavioral dynamics associated with force manipulation (Cavina-Pratesi et al., 2018; Mishra & Thrasher, 2021; Parikh & Santello, 2017; Rao et al., 2021; Rao & Parikh, 2019b; Schettino et al., 2015; Schulz et al., 2016). Within the parietal lobe, the primary sensory, anterior intraparietal sulcus, and the posterior parietal cortical areas have been known to underlie the sensorimotor processing at rest, while executing a reach, or while configuring grasp parameters (Allart et al., 2017; Haar et al., 2017; Rao, Chen, et al., 2020; Tunik et al., 2005). Future research could consider probing the fronto-parietal cortical circuitry to understand how variability within individual neural populations characterize spatiotemporal components of grasping and force dynamics.

## ACKNOWLEDGEMENTS

This work was supported by the University of Houston, Division of Research High Priority Area Research Seed Grant and the National Institutes of Health Eunice Kennedy Shriver National Institute of Child Health and Human Development (NIH/NICHD) R25HD106896 to PJP. The dataset utilized in this study was acquired with support from the National Science Foundation Award IIS-1219231 (J C-V)

## REFERENCES

1. Aguirre, G. K., Zarahn, E., & D’Esposito, M. (1998). The variability of human, BOLD hemodynamic responses. NeuroImage. https://doi.org/10.1006/nimg.1998.0369

2. Alahmadi, A. A. S., Samson, R. S., Gasston, D., Pardini, M., Friston, K. J., D’Angelo, E., Toosy, A. T., & Wheeler-Kingshott, C. A. M. (2016). Complex motor task associated with non-linear BOLD responses in cerebro-cortical areas and cerebellum. Brain Structure and Function, 221(5), 2443–2458. https://doi.org/10.1007/s00429-015-1048-1

3. Allart, E., Delval, A., Caux-Dedeystere, A., Labreuche, J., Viard, R., Lopes, R., & Devanne, H. (2017). Parietomotor connectivity in the contralesional hemisphere after stroke: A paired-pulse TMS study. Clinical Neurophysiology, 128(5), 707– 715. https://doi.org/10.1016/j.clinph.2017.02.016

4. An, J., Yadav, T., Ahmadi, M. B., Tarigoppula, V. S. A., & Francis, J. T. (2018). Near Perfect Neural Critic from Motor Cortical Activity Toward an Autonomously Updating Brain Machine Interface. 2018 40th Annual International Conference of the IEEE Engineering in Medicine and Biology Society (EMBC), 73–76. https://doi.org/10.1109/EMBC.2018.8512274

5. Atique, M. M. U., & Francis, J. T. (2021). Mirror neurons are modulated by grip force and reward expectation in the sensorimotor cortices (S1, M1, PMd, PMv). Scientific Reports, 11(1), 1–17. https://doi.org/10.1038/s41598-021-95536-z

6. Bhagat, N. A., Venkatakrishnan, A., Abibullaev, B., Artz, E. J., Yozbatiran, N., Blank, A. A., French, J., Karmonik, C., Grossman, R. G., O’Malley, M. K., Francisco, G. E., & Contreras-Vidal, J. L. (2016). Design and Optimization of an EEG-Based Brain Machine Interface (BMI) to an Upper-Limb Exoskeleton for Stroke Survivors. Frontiers in Neuroscience, 10, 122. https://doi.org/10.3389/fnins.2016.00122

7. Birn, R. M. (2012). The role of physiological noise in resting-state functional connectivity. NeuroImage, 62(2), 864–870. https://doi.org/10.1016/j.neuroimage.2012.01.016

8. Bradberry, T. J., Gentili, R. J., & Contreras-Vidal, J. L. (2010). Reconstructing three-dimensional hand movements from noninvasive electroencephalographic signals. Journal of Neuroscience, 30(9), 3432–3437. https://doi.org/10.1523/JNEUROSCI.6107-09.2010

9. Bradberry, T. J., Rong, F., & Contreras-Vidal, J. L. (2009). Decoding center-out hand velocity from MEG signals during visuomotor adaptation. NeuroImage, 47(4), 1691–1700. https://doi.org/10.1016/j.neuroimage.2009.06.023

10. Cavina-Pratesi, C., Connolly, J. D., Monaco, S., Figley, T. D., Milner, A. D., Schenk, T., & Culham, J. C. (2018). Human neuroimaging reveals the subcomponents of grasping, reaching and pointing actions. Cortex, 98, 128–148. https://doi.org/10.1016/j.cortex.2017.05.018

11. Chu, W. T. V., & Sanger, T. D. (2009). Force variability during isometric biceps contraction in children with secondary dystonia dueto cerebral palsy. Movement Disorders, 24(9), 1299–1305. https://doi.org/10.1002/mds.22573

12. Churchland, M. M., Santhanam, G., & Shenoy, K. V. (2006). Preparatory Activity in Premotor and Motor Cortex Reflects the Speed of the Upcoming Reach. J Neurophysiol, 96, 3130–3146. https://doi.org/10.1152/jn.00307.2006.

13. Cruz-Garza, J. G., Sujatha Ravindran, A., Kopteva, A. E., Rivera Garza, C., & Contreras-Vidal, J. L. (2020). Characterization of the Stages of Creative Writing With Mobile EEG Using Generalized Partial Directed Coherence. Frontiers in Human Neuroscience, 14, 577651. https://doi.org/10.3389/fnhum.2020.577651

14. Davare, M., Kraskov, A., Rothwell, J. C., & Lemon, R. N. (2011). Interactions between areas of the cortical grasping network. Current Opinion in Neurobiology, 21(4), 565–570. https://doi.org/10.1016/j.conb.2011.05.021

15. Davare, M., Rothwell, J. C., & Lemon, R. N. (2010). Causal Connectivity between the Human Anterior Intraparietal Area and Premotor Cortex during Grasp. Current Biology, 20(2), 176–181. https://doi.org/10.1016/j.cub.2009.11.063

16. de Freitas, P. B., & Lima, K. C. A. (2013). Grip force control during simple manipulation tasks in non-neuropathic diabetic individuals. Clinical Neurophysiology, 124(9), 1904–1910. https://doi.org/10.1016/j.clinph.2013.04.002

17. Delorme, A., & Makeig, S. (2004). EEGLAB: An open source toolbox for analysis of single-trial EEG dynamics including independent component analysis. Journal of Neuroscience Methods, 134(1), 9–21. https://doi.org/10.1016/j.jneumeth.2003.10.009

18. Ehrsson, H. H., Fagergren, A., Johansson, R. S., & Forssberg, H. (2003). Evidence for the involvement of the posterior parietal cortex in coordination of fingertip forces for grasp stability in manipulation. Journal of Neurophysiology, 90(5), 2978–2986.

19. Ehrsson, H. H., Fagergren, A., Jonsson, T., Westling, G., Johansson, R. S., & Forssberg, H. (2000). Cortical Activity in Precision-Versus Power-Grip Tasks: An fMRI Study. Journal of Neurophysiology, 83(1), 528–536. https://doi.org/10.1152/jn.2000.83.1.528

20. Ehrsson, H. H., Fagergren, A., Jonsson, T., Westling, G., Roland, S., Forssberg, H., & Oran, G. (2000). Cortical Activity in Precision-Versus Power-Grip Tasks: An fMRI Study Cortical Activity in Precision-Versus Power-Grip Tasks: An fMRI Study. Journal of Neurophysiology, 83, 528–536.

21. Feeney, D. F., Mani, D., & Enoka, R. M. (2018). Variability in common synaptic input to motor neurons modulates both force steadiness and pegboard time in young and older adults. Journal of Physiology, 596(16), 3793–3806. https://doi.org/10.1113/JP275658

22. Fellows, S. J., & Noth, J. (2003). Grip Force Abnormalities in De Novo Parkinson’s Disease. Movement Disorders, 19(5), 560–565. https://doi.org/10.1002/mds.10695

23. Flanagan, J. R., & Beltzner, M. A. (2000). Independence of perceptual and sensorimotor predictions in the size-weight illusion. Nature Neuroscience, 3(7), 737–741. https://doi.org/10.1038/76701

24. Garrett, D. D., Kovacevic, N., McIntosh, A. R., & Grady, C. L. (2010). Blood oxygen level-dependent signal variability is more than just noise. Journal of Neuroscience, 30(14), 4914–4921. https://doi.org/10.1523/JNEUROSCI.5166-09.2010

25. Garrett, D. D., Kovacevic, N., McIntosh, A. R., & Grady, C. L. (2011). The importance of being variable. Journal of Neuroscience, 31(12), 4496–4503. https://doi.org/10.1523/JNEUROSCI.5641-10.2011

26. Garrett, D. D., Kovacevic, N., McIntosh, A. R., & Grady, C. L. (2013). The modulation of BOLD variability between cognitive states varies by age and processing speed. Cerebral Cortex, 23(3), 684–693. https://doi.org/10.1093/cercor/bhs055

27. Garrett, D. D., McIntosh, A. R., & Grady, C. L. (2014). Brain signal variability is parametrically modifiable. Cerebral Cortex, 24(11), 2931–2940. https://doi.org/10.1093/cercor/bht150

28. Garrett, D. D., Samanez-Larkin, G. R., MacDonald, S. W. S., Lindenberger, U., McIntosh, A. R., & Grady, C. L. (2013). Moment-to-moment brain signal variability: A next frontier in human brain mapping? Neuroscience and Biobehavioral Reviews, 37(4), 610–624. https://doi.org/10.1016/j.neubiorev.2013.02.015

29. Goel, R., Nakagome, S., Rao, N., Contreras-Vidal, J., & Parikh, P. (2017). Role of supplementary motor area in postural control. Neuroscience (Annual Meeting of the Society for Neuroscience).

30. Goel, R., Nakagome, S., Rao, N., Paloski, W. H., Contreras-Vidal, J. L., & Parikh, P. J. (2019). Fronto-Parietal Brain Areas Contribute to the Online Control of Posture during a Continuous Balance Task. Neuroscience, 413, 135–153. https://doi.org/10.1016/j.neuroscience.2019.05.063

31. Grady, C. L., & Garrett, D. D. (2014). Understanding variability in the BOLD signal and why it matters for aging. Brain Imaging and Behavior, 8(2), 274–283. https://doi.org/10.1007/s11682-013-9253-0

32. Grady, C. L., & Garrett, D. D. (2018). Brain signal variability is modulated as a function of internal and external demand in younger and older adults. NeuroImage, 169(December 2017), 510–523. https://doi.org/10.1016/j.neuroimage.2017.12.031

33. Grafton, S. T. (2010). The cognitive neuroscience of prehension: Recent developments. Experimental Brain Research, 204(4), 475–491. https://doi.org/10.1007/s00221-010-2315-2

34. Haar, S., Donchin, O., & Dinstein, I. (2017). Individual Movement Variability Magnitudes Are Explained by Cortical Neural Variability. The Journal of Neuroscience, 37(37), 9076–9085. https://doi.org/10.1523/JNEUROSCI.1650-17.2017

35. Hendrix, C. M., Mason, C. R., & Ebner, T. J. (2009). Signaling of Grasp Dimension and Grasp Force in Dorsal Premotor Cortex and Primary Motor Cortex Neurons During Reach to Grasp in the Monkey. Journal of Neurophysiology, 102(1), 132–145. https://doi.org/10.1152/jn.00016.2009

36. Kilicarslan, A., Grossman, R. G., & Contreras-Vidal, J. L. (2016). A robust adaptive denoising framework for real-time artifact removal in scalp EEG measurements. Journal of Neural Engineering, 13(2), 026013. https://doi.org/10.1088/1741-2560/13/2/026013

37. Kilner, J. M., Salenius, S., Baker, S. N., Jackson, A., Hari, R., & Lemon, R. N. (2003). Task-dependent modulations of cortical oscillatory activity in human subjects during a bimanual precision grip task. NeuroImage, 18(1), 67–73. https://doi.org/10.1006/nimg.2002.1322

38. Kontson, K. L., Megjhani, M., Brantley, J. A., Cruz-Garza, J. G., Nakagome, S., Robleto, D., White, M., Civillico, E., & Contreras-Vidal, J. L. (2015). Your Brain on Art: Emergent Cortical Dynamics During Aesthetic Experiences. Frontiers in Human Neuroscience, 9, 626. https://doi.org/10.3389/fnhum.2015.00626

39. Kumar, D., Das, A., Lahiri, U., & Dutta, A. (2016). A human-machine-interface integrating low-cost sensors with a neuromuscular electrical stimulation system for post-stroke balance rehabilitation. Journal of Visualized Experiments, 110(52394). https://doi.org/10.3791/52394

40. Kumar, D., Verma, S., Bhattacharya, S., & Lahiri, U. (2016). Audio-Visual Stimulation in Conjunction with Functional Electrical Stimulation to Address Upper Limb and Lower Limb Movement Disorder. European Journal of Translational Myology, 26(2), 6030. https://doi.org/10.4081/ejtm.2016.6030

41. Lodha, N., & Christou, E. A. (2017). Low-frequency oscillations and control of the motor output. Frontiers in Physiology, 8(FEB), 1–9. https://doi.org/10.3389/fphys.2017.00078

42. Lodha, N., Misra, G., Coombes, S. A., Christou, E. A., & Cauraugh, J. H. (2013).

43. Increased force variability in chronic stroke: Contributions of force modulation below 1 Hz. PLoS ONE, 8(12), 1–9. https://doi.org/10.1371/journal.pone.0083468

44. Lukos, J. R., Cho, J. Y., & Santello, M. (2013). Grasping uncertainty: Effects of sensorimotor memories on high-level planning of dexterous manipulation. Journal of Neurophysiology, 109(12), 2937–2946. https://doi.org/10.1152/jn.00060.2013

45. Luu, T. P., He, Y., Brown, S., Nakagame, S., & Contreras-Vidal, J. L. (2016). Gait adaptation to visual kinematic perturbations using a real-time closed-loop brain– computer interface to a virtual reality avatar. Journal of Neural Engineering, 13(3), 036006. https://doi.org/10.1088/1741-2560/13/3/036006

46. Luu, T. P., Nakagome, S., He, Y., & Contreras-Vidal, J. L. (2017). Real-time EEG-based brain-computer interface to a virtual avatar enhances cortical involvement in human treadmill walking. Scientific Reports, 7(1), 1–12. https://doi.org/10.1038/s41598-017-09187-0

47. Mcintosh, A. R., Kovacevic, N., & Itier, R. J. (2008). Increased Brain Signal Variability Accompanies Lower Behavioral Variability in Development. 4(7). https://doi.org/10.1371/journal.pcbi.1000106

48. McIntosh, A. R., Kovacevic, N., Lippe, S., Garrett, D., Grady, C., & Jirsa, V. (2010). The development of a noisy brain. Archives Italiennes de Biologie, 148(3), 323–337. https://doi.org/10.4449/aib.v148i3.1225

49. Mishra, R. K., Park, C., Zhou, H., Najafi, B., & Thrasher, T. A. (2022). Evaluation of Motor and Cognitive Performance in People with Parkinson’s Disease Using Instrumented Trail-Making Test. Gerontology, 68(2), 234–240. https://doi.org/10.1159/000515940

50. Mishra, R. K., & Thrasher, A. T. (2021). Transcranial direct current stimulation of dorsolateral prefrontal cortex improves dual-task gait performance in patients with Parkinson’s disease: A double blind, sham-controlled study. Gait & Posture, 84, 11–16. https://doi.org/10.1016/j.gaitpost.2020.11.012

51. Moore, B., Khang, S., & Francis, J. T. (2020). Noise-Correlation Is Modulated by Reward Expectation in the Primary Motor Cortex Bilaterally During Manual and Observational Tasks in Primates. Frontiers in Behavioral Neuroscience, 14(December), 1–15. https://doi.org/10.3389/fnbeh.2020.541920

52. Olivier, E., Davare, M., Andres, M., & Fadiga, L. (2007). Precision grasping in humans: From motor control to cognition. Current Opinion in Neurobiology, 17(6), 644– 648. https://doi.org/10.1016/j.conb.2008.01.008

53. Paek, A. Y., Gailey, A., Parikh, P. J., Santello, M., & Contreras-Vidal, J. L. (2019). Regression-based reconstruction of human grip force trajectories with noninvasive scalp electroencephalography. Journal of Neural Engineering, 16(6), 66030. https://doi.org/10.1088/1741-2552/ab4063

54. Parikh, P. J., & Cole, K. J. (2015). Effects of Transcranial Direct Current Stimulation on the Control of Finger Force during Dexterous Manipulation in Healthy Older Adults. Plos One, 10(4), e0124137. https://doi.org/10.1371/journal.pone.0124137

55. Parikh, P. J., Fine, J. M., & Santello, M. (2020). Dexterous Object Manipulation Requires Context-Dependent Sensorimotor Cortical Interactions in Humans. Cerebral Cortex, 30(5), 3087–3101. https://doi.org/10.1093/cercor/bhz296

56. Parikh, P. J., & Santello, M. (2017). Role of human premotor dorsal region in learning a conditional visuomotor task. Journal of Neurophysiology, 117(1), 445–456. https://doi.org/10.1152/jn.00658.2016

57. Park, C., Mishra, R., Sharafkhaneh, A., Bryant, M. S., Nguyen, C., Torres, I., Naik, A. D., & Najafi, B. (2021). Digital Biomarker Representing Frailty Phenotypes: The Use of Machine Learning and Sensor-Based Sit-to-Stand Test. Sensors (Basel, Switzerland), 21(9). https://doi.org/10.3390/s21093258

58. Perez, M. A., & Cohen, L. G. (2009). Scaling of motor cortical excitability during unimanual force generation. Cortex; a Journal Devoted to the Study of the Nervous System and Behavior, 45(9), 1065–1071. https://doi.org/10.1016/j.cortex.2008.12.006

59. Poon, C., Chin-Cottongim, L. G., Coombes, S. A., Corcos, D. M., & Vaillancourt, D. E. (2012). Spatiotemporal dynamics of brain activity during the transition from visually guided to memory-guided force control. Journal of Neurophysiology, 108(5), 1335–1348. https://doi.org/10.1152/jn.00972.2011

60. Poon, C., Coombes, S. A., Corcos, D. M., Christou, E. A., & Vaillancourt, D. E. (2013). Transient shifts in frontal and parietal circuits scale with enhanced visual feedback and changes in force variability and error. Journal of Neurophysiology, 109(8), 2205–2215. https://doi.org/10.1152/jn.00969.2012

61. Rao, N., Chen, Y. T., Ramirez, R., Tran, J., Li, S., & Parikh, P. J. (2020). Time-course of pain threshold after continuous theta burst stimulation of primary somatosensory cortex in pain-free subjects. Neuroscience Letters, 722, 134760. https://doi.org/10.1016/j.neulet.2020.134760

62. Rao, N., Chen, Y.-T., Ramirez, R., Tran, J., Li, S., & Parikh, P. J. (2019). Persistent Elevation of Electrical Pain Threshold following Continuous Theta Burst Stimulation over Primary Somatosensory Cortex in Humans. BioRxiv, 724344.

63. Rao, N., Mehta, N., Patel, P., & Parikh, P. J. (2020). Modulation of Grasp Parameters using Arbitrary Cues about Object Property in Older Adults. BioRxiv. https://doi.org/10.1101/2020.10.19.344457

64. Rao, N., Mehta, N., Patel, P., & Parikh, P. J. (2021). Effects of aging on conditional visuomotor learning for grasping and lifting eccentrically weighted objects.

65. Journal of Applied Physiology, 131(3), 937–948. https://doi.org/10.1152/japplphysiol.00932.2020

66. Rao, N., & Parikh, P. J. (2017). Variability in Corticospinal Excitability during Digit Force Planning for Grasping in Humans. Neuroscience (Annual Meeting of the Society for Neuroscience).

67. Rao, N., & Parikh, P. J. (2019a). Fluctuations in Human Corticospinal Activity Prior to Grasp. Frontiers in Systems Neuroscience, 13(December), 1–15. https://doi.org/10.3389/fnsys.2019.00077

68. Rao, N., & Parikh, P. J. (2019b). Intertrial Variability in Human Corticospinal Activity during Grasp Force Planning. BioRxiv, 676833. https://doi.org/10.1101/676833

69. Rao, N., Skinner, L., Kass, J., & Parikh, P. J. (2019). Contribution of Human Primary Motor Cortex to Force Variability during Precision Grasping. Neuroscience (Annual Meeting of the Society for Neuroscience).

70. Rastogi, A., Vargas-Irwin, C. E., Willett, F. R., Abreu, J., Crowder, D. C., Murphy, B. A., Memberg, W. D., Miller, J. P., Sweet, J. A., Walter, B. L., Cash, S. S., Rezaii, P. G., Franco, B., Saab, J., Stavisky, S. D., Shenoy, K. V., Henderson, J. M., Hochberg, L. R., Kirsch, R. F., & Ajiboye, A. B. (2020). Neural Representation of Observed, Imagined, and Attempted Grasping Force in Motor Cortex of Individuals with Chronic Tetraplegia. Scientific Reports, 10(1), 1–16. https://doi.org/10.1038/s41598-020-58097-1

71. Ravindran, A. S., Malaya, C. A., John, I., Francisco, G. E., Layne, C., & Contreras-Vidal, J. L. (2022). Decoding neural activity preceding balance loss during standing with a lower-limb exoskeleton using an interpretable deep learning model. Journal of Neural Engineering, 19(3), 36015. https://doi.org/10.1088/1741-2552/ac6ca9

72. Sanchez, J. C., Gunduz, A., Carney, P. R., & Principe, J. C. (2008). Extraction and localization of mesoscopic motor control signals for human ECoG neuroprosthetics. Journal of Neuroscience Methods, 167(1), 63–81. https://doi.org/10.1016/j.jneumeth.2007.04.019

73. Schalk, G., Kubánek, J., Miller, K. J., Anderson, N. R., Leuthardt, E. C., Ojemann, J. G., Limbrick, D., Moran, D., Gerhardt, L. A., & Wolpaw, J. R. (2007). Decoding two-dimensional movement trajectories using electrocorticographic signals in humans. Journal of Neural Engineering, 4(3), 264–275. https://doi.org/10.1088/1741-2560/4/3/012

74. Schettino, L. F., Adamovich, S. V., Bagce, H., Yarossi, M., & Tunik, E. (2015). Disruption of activity in the ventral premotor but not the anterior intraparietal area interferes with on-line correction to a haptic perturbation during grasping. The Journal of Neuroscience : The Official Journal of the Society for Neuroscience, 35(5), 2112–2117. https://doi.org/10.1523/JNEUROSCI.3000-14.2015

75. Schulz, R., Buchholz, A., Frey, B. M., Bönstrup, M., Cheng, B., Thomalla, G., Hummel, F. C., & Gerloff, C. (2016). Enhanced Effective Connectivity between Primary Motor Cortex and Intraparietal Sulcus in Well-Recovered Stroke Patients. Stroke, 47(2), 482–489. https://doi.org/10.1161/STROKEAHA.115.011641

76. Shah-Basak, P. P., Sivaratnam, G., Teti, S., Francois-Nienaber, A., Yossofzai, M., Armstrong, S., Nayar, S., Jokel, R., & Meltzer, J. (2020). High definition transcranial direct current stimulation modulates abnormal neurophysiological activity in post-stroke aphasia. Scientific Reports, 10(1), 1–18. https://doi.org/10.1038/s41598-020-76533-0

77. Smith, B. W., Rowe, J. B., & Reinkensmeyer, D. J. (2018). Real-time slacking as a default mode of grip force control: Implications for force minimization and personal grip force variation. Journal of Neurophysiology, 120(4), 2107–2120. https://doi.org/10.1152/jn.00700.2017

78. Svendsen, J. H., & Madeleine, P. (2010). Amount and structure of force variability during short, ramp and sustained contractions in males and females. Human Movement Science, 29(1), 35–47. https://doi.org/10.1016/j.humov.2009.09.001

79. Tunik, E., Frey, S. H., & Grafton, S. T. (2005). Virtual lesions of the anterior intraparietal area disrupt goal-dependent on-line adjustments of grasp. Nature Neuroscience, 8(4), 505–511. https://doi.org/10.1038/nn1430

80. Vaillancourt, D. E., Thulborn, K. R., & Corcos, D. M. (2003). Neural Basis for the Processes That Underlie Visually Guided and Internally Guided Force Control in Humans. Journal of Neurophysiology, 90(5), 3330–3340. https://doi.org/10.1152/jn.00394.2003

81. van Polanen, V., & Davare, M. (2015). Interactions between dorsal and ventral streams for controlling skilled grasp. Neuropsychologia, 79, 186–191. https://doi.org/10.1016/j.neuropsychologia.2015.07.010

82. Vieluf, S., Temprado, J., Berton, E., Jirsa, V. K., & Sleimen-malkoun, R. (2015). Effects of task and age on the magnitude and structure of force fluctuations: Insights into underlying neuro-behavioral processes. 1–17. https://doi.org/10.1186/s12868-015-0153-7

83. Yadav, T., Uddin Atique, M. M., Fekri Azgomi, H., Francis, J. T., & Faghih, R. T. (2019). Emotional Valence Tracking and Classification via State-Space Analysis of Facial Electromyography. 2019 53rd Asilomar Conference on Signals, Systems, and Computers, 2116–2120. https://doi.org/10.1109/IEEECONF44664.2019.9048868

84. Young, D. R., Parikh, P. J., & Layne, C. S. (2020). Non-invasive Brain Stimulation of the Posterior Parietal Cortex Alters Postural Adaptation. Frontiers in Human Neuroscience, 14(June), 1–10. https://doi.org/10.3389/fnhum.2020.00248

